# The P387 Thrombospondin-4 Variant Promotes Accumulation of Macrophages in Atherosclerotic Lesions

**DOI:** 10.1101/666602

**Authors:** Santoshi Muppala, Mohammed Tanjimur Rahman, Irene Krukovets, Dmitriy Verbovetskiy, Elzbieta Pluskota, Aaron Fleischman, D. Geoffrey Vince, Edward F. Plow, Olga Stenina-Adognravi

## Abstract

**Aims:** Thrombopspondin-4 (TSP4) is a pro-angiogenic protein that has been implicated in tissue remodeling and local vascular inflammation. TSP4 and, in particular, its SNP variant, P387 TSP4, have been associated with cardiovascular disease.

Macrophages are central to initiation and resolution of inflammation and development of atherosclerotic lesions, but the effects of the P387 TSP4 on macrophages remain essentially unknown. We examined the effects of the P387 TSP4 variant on macrophages in cell culture and *in vivo* in a murine model of atherosclerosis. Further, the levels and distributions of the twoTSP4 variants were assessed in human atherosclerotic arteries.

**Methods and Results:** In *ApoE*^−/−^/P387-TSP4 knock-in mice, atherosclerotic lesions accumulated more macrophages than lesions bearing A387 TSP4. The levels of inflammatory markers were increased in lesions of *ApoE*^−/−^/P387-TSP4 knock-in mice compared to *ApoE*^−/−^ mice. Lesions in human arteries from individuals carrying the P387 variant had higher levels of TSP4 and higher macrophage accumulation. P387 TSP4 was more active in supporting adhesion of cultured human and mouse macrophages in experiments using recombinant TSP4 variants and in cells derived from P387-TSP4 knock-in mice.

**Conclusions:** TSP4 supports the adhesion of macrophages and their accumulation in atherosclerotic lesions. P387 TSP4 is more active in supporting these pro-inflammatory events in the vascular wall, which may contribute to the increased association of P387 TSP4 with cardiovascular disease.

**Abbreviations:** BSA, bovine serum albumin; DMSO, dimethyl sulfoxide; ECM, extracellular matrix; *Thbs4*^−/−^, thrombospondin-4 gene knock-out; WT, wild type; P387-TSP4 KI, P387 *TSP4* knock-in mice; OCT, Optimum Cutting Temperature; vWF, von Willebrand factor; α-SMA, alpha-smooth muscle actin; Egr2, Early Growth Response 2; PBS, Phosphate Buffer saline; DMEM, Dulbecco’s Modified Eagle Medium.

## 1. Introduction

Thrombospondin-4 (TSP4) is a matricellular protein from the thrombospondin (TSP) family, which consists of five homologous but differentially expressed members that are products of five different genes on different chromosomes (1, 2). Recent studies have demonstrated the importance of TSP4 in the cardiovascular system (3–17), cancer (15, 18–27), and the nervous system (28–32). These reports have placed considerable attention on this poorly studied TSP, but many of its functions and its regulation still remain unknown. Our previous work in *Thbs4*^−/−^ mice (17) revealed that TSP4 promotes atherosclerotic lesions, regulates the local vascular inflammation, and supports accumulation of macrophages in atherosclerotic lesion.

The effects of TSP4 depend on its interaction with cellular receptors and ligands in the extracellular matrix (ECM). P387-TSP4, a SNP variant common in North American population (3, 6) has been associated with myocardial infarction (MI) and coronary artery disease (CAD) (3–10). Previously, we reported that P387-TSP4 is less susceptible to proteolysis and is more active in regulating cellular functions (14, 33–35). Here, we used P387-TSP4 knock-in (KI) mice and recombinant P387-TSP4 to examine the activity of this variant in atherosclerosis. We compared the effects of TSP4 variants not only in mouse but also in human atherosclerotic arteries and identified both commonalities and differences in the effects of the TSP4 variants on macrophages.

## 2. Materials and Methods

### 2.1. Animals

Mice were in a C57BL/6 background. Both genders were used, but no gender-specific differences were detected. *Thbs4*^−/−^ mice were described previously (14–17, 36). Control wild-type (WT) mice were from the same mouse colony as *Thbs4*^−/−^ mice or P387-TSP4-KI mice. For experiments exclusively using WT mice, WT C57BL/6 mice were purchased from the Jackson Laboratories. Animals were housed in the AAALAC-approved animal facilities of the Cleveland Clinic. Animal procedures were approved by the Institutional Animal Care and Use Committee of the Cleveland Clinic and were in accord with the NIH Guide for Animal Use. Ketamine (100 mg/kg)/Xylazine (15 mg/kg) was used for anesthesia.

### 2.2. Atherosclerosis model and analysis of the lesions

For the analysis of atherosclerotic lesions, four groups of mice (n ≥ 13) were analyzed; *ApoE*^−/−^ and P387-TSP4-KI/*ApoE*^−/−^ mice were maintained either on a regular chow or a Western diet (TD88137, Teklad Diets, Madison, WI). All mice were analyzed for development of atherosclerosis at 20 weeks of age, 16 weeks after being placed on the two diets. In mice on both diets, lesions that developed in the aortic root, aortic arch, descending aorta and femoral artery were measured. At 20 weeks of age, the circulatory system of anesthetized mice was perfused with 0.9% NaCl by cardiac intraventricular cannulation. Surface lesion area was determined by an *en face* method. Oil Red O-stained areas of lesions were quantified using ImagePro 6.3.

### 2.3. Genotyping

Genomic DNAs were extracted from frozen tissue samples using Qiagen DNeasy kit. TSP4 A387P mutations were detected using rhAmp™ SNP Assay (SNP id: rs1866389, Integrated DNA Technologies) kit with Quantstudio 5 cycler genotyping program according to assay protocol set by the manufacturer. Genotyping calls were made by the automated program with standards (WT and A387P variant) set with gBlocks™ Gene Fragments (Integrated DNA Technologies) as suggested by manufacturer’s instructions for the kit.

### 2.4. Human Coronary Artery Samples

A total of 24 fresh human coronary artery samples were obtained from the Histochemistry Core of the Department of Biomedical Engineering of the Cleveland Clinic, embedded in Tissue-Tek® O.C.T. Compound (#4583, Sakura Finetek) followed by flash freeze in liquid nitrogen. Frozen tissue section, 10 μm thickness, were cut with microtome and mounted on slides for immuno-histochemical or immunofluorescence staining.

### 2.5. Blood cell count

200 μl of blood was collected from heart into 0.6 ml snap cap aliquot tubes. EDTA coated syringes were used. Samples were analyzed within one hour of collection. Cells were counted in the Laboratory Diagnostics Core at the Cleveland Clinic using Advia hematology analyzer.

### 2.6. Human blood-derived monocyte isolation

Blood monocytes were isolated using Ficoll Paque, and total number of cells were counted.

### 2.7. Cell Culture and Cell Lines

Human macrophage-like cell line WBC264-9C were purchased from ATCC (Virginia, USA). Eagle’s Minimum Essential Medium (WBC264-9C), supplemented with 10% fetal bovine serum (FBS), sodium bi-carbonate (1.5gm/L), 100 U/mL penicillin, and 100 µg/mL streptomycin, was used for culture at 37 °C in 5% CO_2_ atmosphere according to ATCC recommendations. All experiments were done with cells between passages 4 and 8.

### 2.8. Isolation of Peritoneal Mouse Macrophages

Mouse peritoneal macrophages (MPM) were isolated from WT, *Thbs4*^−/−^ and P387-TSP4-KI mice and directly used for adhesion and migration experiments. MPM were derived from mice treated with LPS (0.25 μg/g) for 72 hours by peritoneal lavage with PBS.

Initial attachment of cells was determined after 3 hours using CyQUANT live cell quantification kits. Each group of live cells was normalized to its respective 3 hours initial attachment to account for the differences in initial adhesion of macrophages.

### 2.9. Purification of recombinant TSP4 variants

Recombinant TSP4 (rTSP4) variants were purified from culture supernatants of HEK293 cells stably transfected with A387- and P387-coding cDNA of TSP4 as was described in our earlier publications (27, 33, 34).

### 2.10. In vitro adhesion assays

Adhesion assays were performed as previously described (14, 15, 17, 33, 35). For human blood monocytes, 1×10^5^ cells/well in 100 μl of DMEM-F-12 medium containing 10% FBS were added to 96-well TC-nontreated plates (Becton Dickinson, Franklin Lakes, NJ) coated with immobilized purified recombinant P387 TSP4 or A387 TSP4. The wells were coated with 100 μl P387 TSP4 or A387 TSP4 (10 μg/ml) for 2 hours at 37ºC, post-coated with 0.5% polyvinylpyrrolidone for 1 hour at 37ºC, washed twice with PBS, and incubated for 0.5 or 4h at 37ºC (33–36). Non-adherent cells were aspirated and 150 μl of Hanks balanced saline solution containing 1 mM Mg^2+^, 1 mM Ca^2+^ and 0.1% bovine serum albumin (BSA) were added to each well.

### 2.11. H_2_O_2_ release

H_2_O_2_ was measured in the supernatants of adherent human monocytes after 48 hours using the Amplex Red Hydrogen Peroxide/Peroxidase Assay Kit (Thermofisher Scientific, Grand Island, NY), according to manufacturer’s instructions.

### 2.12. In vitro migration assays

Cells were suspended in the serum-free DMEM and transferred into the trans-well chambers (Corning, Corning, NY, USA). FBS (20%) was used as a chemo-attractant in the bottom chambers. The cells were incubated at 37 °C for 4 hr, the medium was aspirated, and attached cells were removed from the surface of the upper chamber using Q-tips. The plates were frozen at − 80 °C for 3 hours, and DNA of remaining cells was quantified using CyQUANT reagent (Invitrogen, Carlsbad, CA, USA).

### 2.13. Immunohistochemistry, immunofluorescence, confocal imaging and quantification of macrophage markers

Sections of aortic lesions were stained with primary rat anti-mouse CD68 (MCA1957B, Bio-Rad), rabbit anti-mouse CD38 (bs-0979R, Bioss Antibodies), rabbit anti-mouse Egr2 (ab90518, Abcam), rabbit anti-mouse MCP-1 (Abcam), or rabbit anti-mouse NF-kB p65 (phospho S536, Abcam) antibody. For human arteries staining, rabbit anti-human vWF antibody (A0082, Dako), mouse anti-human CD68 (Dako) and goat anti-humanTSP4 (AF2390, R&D Systems) (1:200) were used followed by the treatment with Vecta Stain ABC Kit. Visualization after staining with the antibodies was performed using a high-resolution Aperio AT2 slide scanner (Leica Microsystems, GmbH, Wetzlar, Germany). Images of whole slides at 20x magnification were scanned, and the staining was quantified to determine the percentage of stained area using Image Pro and Adobe Photoshop CS6 (Media Cybernetics). The person performing quantification was blinded to the assignment of animals between groups. For immunofluorescence, rat anti-mouse CD68 (MCA1957B, Bio-Rad), goat anti-human TSP4 (AF2390, R&D systems), monoclonal mouse anti-human alpha smooth muscle actin (M0851, Dako), monoclonal mouse anti-human CD68 (M0814, Dako), Rabbit anti-human vWF antibody (A0082, Dako) (1:200), were used with corresponding secondary antibodies (1:1000). Secondary antibodies were anti-goat NL557 conjugated donkey IgG (NL001 R&D systems), goat polyclonal antibody to rat IgG Alexa Fluor 488 (ab150161, Abcam), anti-mouse IgG NL493 conjugated donkey IgG (NL009, R&D systems) and Alexa Fluor™647 F(ab’)2 fragment of goat anti-mouse IgG(H+L) (A21237, Invitrogen) and Alexa Fluor™647 chicken anti-rabbit IgG (H+L) (A21443, Invitrogen). Images were taken at a high resolution confocal microscope (Leica DM 2500) at 63x magnification. All sections with primary antibodies were incubated for 2 hours at 4°C followed by incubating sections in secondary antibodies for 45 min at 4°C.

### 2.14. Statistical analysis

Group size was calculated based on the previous data obtained in mouse models (14, 16, 17, 36). Analyses of the data were performed using Sigma Plot Software (Systat Software, San Jose, CA, USA): Student’s t-test and one-way ANOVA were used to determine the significance of parametric data, and Wilcoxon rank sum test was used for nonparametric data. The significance level was set at p=0.05. The data are presented as mean ± S.E.M. Numbers of biological repeats are listed in each legend.

## 3. Results

### 3.1. P387 TSP4 promotes accumulation of macrophages in atherosclerotic lesions of ApoE^−/−^ mice

*ApoE*^*−/−*^(A387) and *ApoE*^*−/−*^/ P387-TSP4-KI mice were fed with either chow or western diet for 16 weeks, starting at 4 weeks of age. Atherosclerotic lesions in aortic arch, descending aorta, and femoral artery of the two genotypes were stained with Oil Red O and quantified as described in Methods. There was no difference in the size of lesions between the two genotypes on either diet (Fig.1A shows the results for mice on Western diet). However, when the cellular content of the lesions was evaluated, the lesions of *ApoE*^*−/−*^/ P387-TSP4-KI mice were found to contain significantly more cells than the lesions of *ApoE*^*−/−*^mice. Immunostaining with anti-CD68 (marker of macrophage) and anti-α-smooth muscle actin (α-SMA) revealed that the number of both cell types was higher in *ApoE*^*−/−*^/ P387-TSP4-KI mice (Fig.1B).

**Figure 1:**
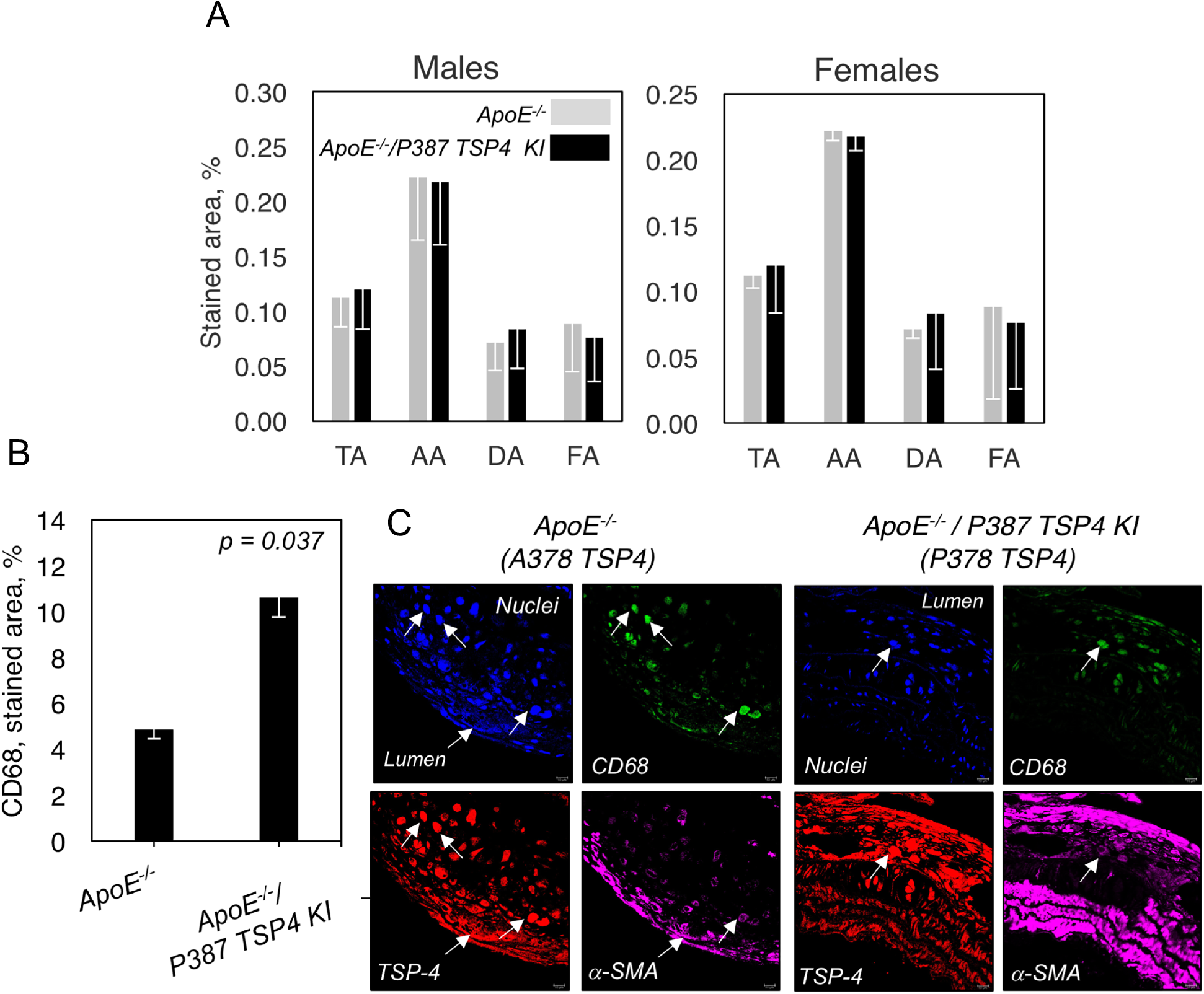
P387 TSP4 promotes accumulation of macrophages in atherosclerotic lesions without changing the accumulation of lipids. **A:** P387 TSP4 does not change the lesion size. TA, total lesion; AA, aortic arch lesion; DA, descending aorta lesion; FA, femoral artery lesion. Western diet, Oil red O, stained area (% of total area), n=14. **B:** Increased accumulation of macrophages in the lesion of *ApoE*^*−/−*^/P387 TSP4 KI mice. Quantification of anti-CD68 (macrophage marker) staining in atherosclerotic lesions of the in *ApoE*^*−/−*^ and *ApoE*^*−/−*^/P387 TSP4 KI mice aorta (n=9). Immunofluorescence, anti-CD68 (green), anti-TSP4 (red), anti-α-SMA, (magenta), DAPI (blue, nuclei). Arrows indicate cells producing TSP4.

Blood cell counts were performed in *Thbs4*^*−/−*^ and P387-TSP4-KI mice to determine whether the higher accumulation of macrophages in lesions may reflect the higher numbers of leukocytes and/or monocytes in blood. There was no difference between the leukocyte, monocyte, or lymphocyte numbers in P387-TSP4-KI and the corresponding WT control mice (Table 1). In *Thbs4*^*−/−*^ mice that had lower macrophages accumulation in lesions (17), the number of leukocytes was significantly higher compared to corresponding control WT mice. There was no significant difference in monocytes numbers, and the difference in leukocyte numbers was attributable to a significant difference in lymphocytes counts.

**Table 1.**
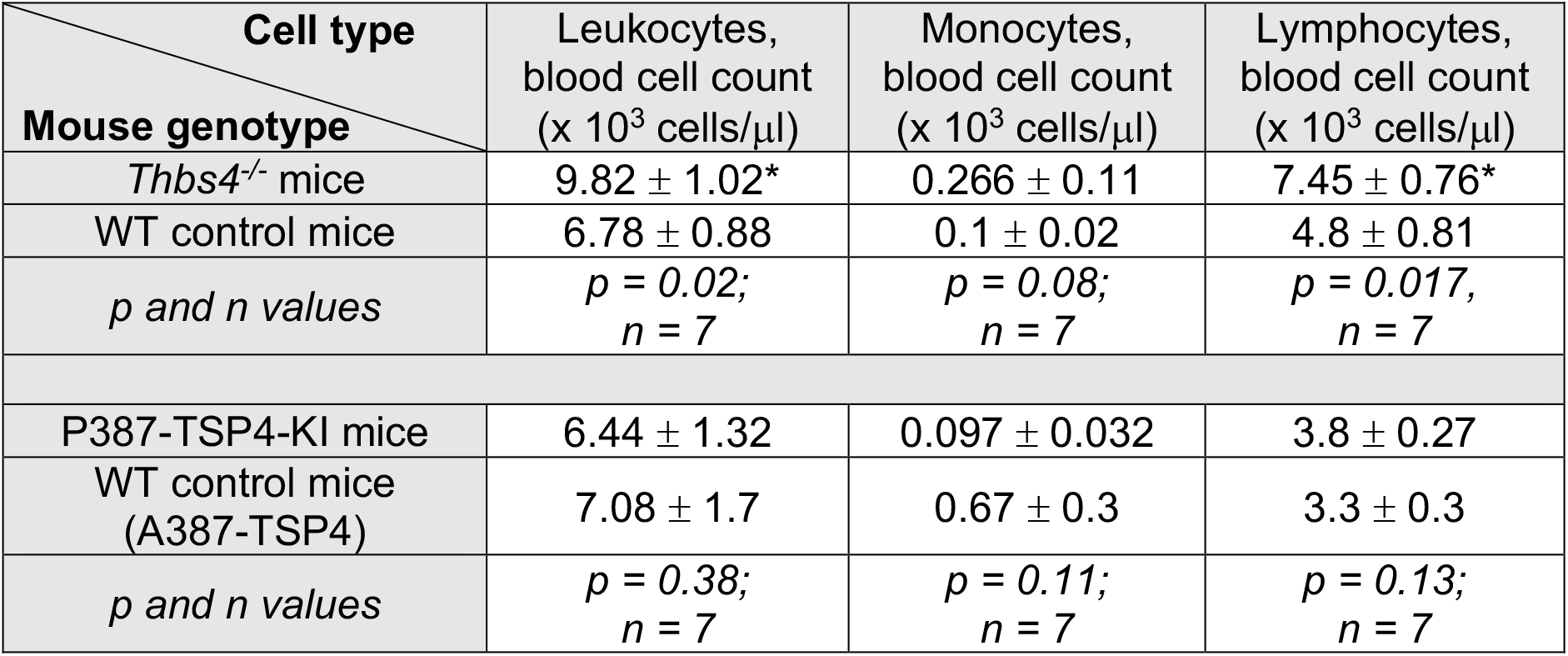
Blood cell counts in Thbs4^*−/−*^ mice, P387-TSP4 KI mice, and corresponding control WT mice (A387). Cell numbers were analyzed in blood as described in Methods.

### 3.2. Levels of CD38, a marker of pro-inflammatory macrophages, are increased in lesions of ApoE^−/−/^ P387-TSP4-KI mice

Co-localization of CD68, a marker of macrophages, and CD38, a marker of pro-inflammatory immune cells was examined in atherosclerotic lesions of mice of both genotypes using immunofluorescence and confocal microscopy (Fig.2A). CD38 staining was associated with macrophages in the lesions. Presence of Egr2, a marker of tissue repair macrophages, was also analyzed (Fig.2B). Egr2 expression was found in macrophages as well as in other cell types, especially in smooth muscle cells.

**Figure 2.**
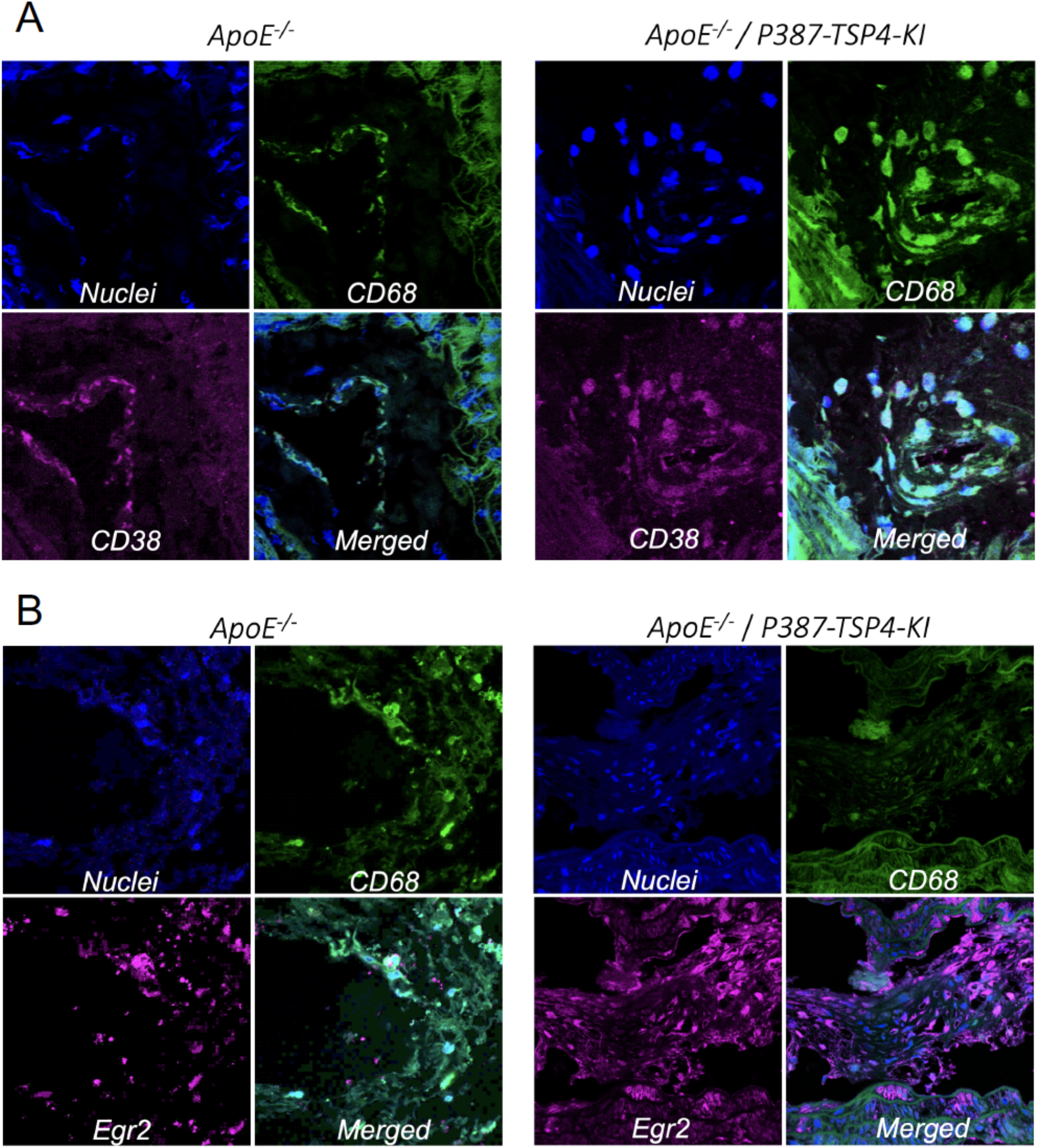
CD38 and Egr2 in lesions. **A:** Atherosclerotic lesions in Aortic Arch of the in *ApoE*^*−/−*^ and *ApoE*^*−/−*^/P387 TSP4 KI mice aorta were stained with pro-inflammatory CD68 (green) and pro-resolving CD38 (magenta) markers were stained with corresponding antibodies as described in Methods. **B:** CD68 (green) and Egr2 (magenta). Immunofluoresecnce, confocal microscopy.

The accumulation of pro-inflammatory macrophages was increased in *ApoE*^*−/−*^/ P387-TSP4-KI mice compared to *ApoE*^*−/−*^ mice expressing A387 TSP4: the level of CD38, a marker of pro-inflammatory macrophages, was significantly increased (Fig.3A, p = 0.007). The level of a marker of tissue repair macrophages, Egr2, was decreased, although this decrease did not reach statistical significance with n = 10 (Fig. 3B, p = 0.08).

**Figure 3.**
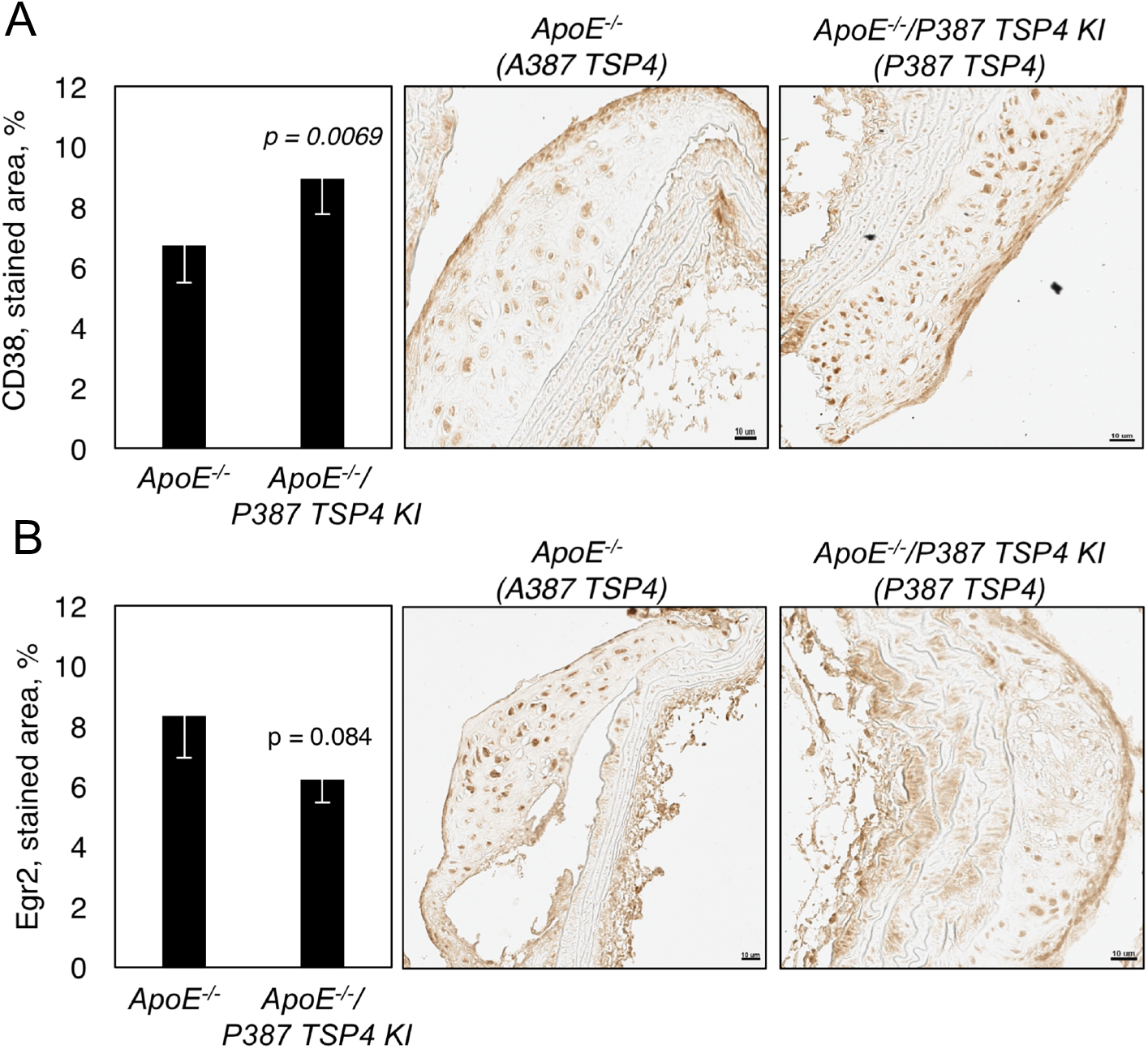
Levels of CD38 (marker of pro-inflammatory macrophages) is increased in atherosclerotic lesions of *ApoE*^*−/−*^/P387 TSP4 KI mice. **A:** Accumulation of pro-inflammatory macrophages is increased in the lesion of *ApoE*^*−/−*^/P387 TSP4 KI mice. CD38 staining in sections of whole atherosclerotic lesions in Aortic Arch in *ApoE*^*−/−*^ and *ApoE*^*−*/−^/P387 TSP4 KI mice aorta (n=10), immunohistochemistry with anti-CD38 antibody. **B:** Accumulation of tissue repair macrophages are decreased in the lesion of *ApoE^−/−^*/P387 TSP4 KI mice. Egr2 staining in sections of atherosclerotic lesions in *ApoE*^*−/−*^ and *ApoE*^*−/−*^/P387 TSP4 KI mice aorta (n=10), immunohistochemistry with anti-Egr2 antibody.

### 3.3. Increased levels of inflammatory markers in lesions of ApoE^−/−^/ P387-TSP4-KI mice

The levels of MCP-1, p65, and ICAM-1 were visualized in lesions of *ApoE*^*−/−*^/ P387-TSP4-KI mice and *ApoE*^*−/−*^ mice expressing A387 TSP4 by immunohistochemistry with anti-MCP-1, anti-p65, and anti-ICAM-1 antibodies as described in Methods. The percentage of stained area of lesions was quantified (Fig.4). The levels of MCP-1, p65, and ICAM-1 were significantly higher in *ApoE*^*−/−*^/ P387-TSP4-KI mice than in *ApoE*^*−/−*^ mice.

**Figure 4.**
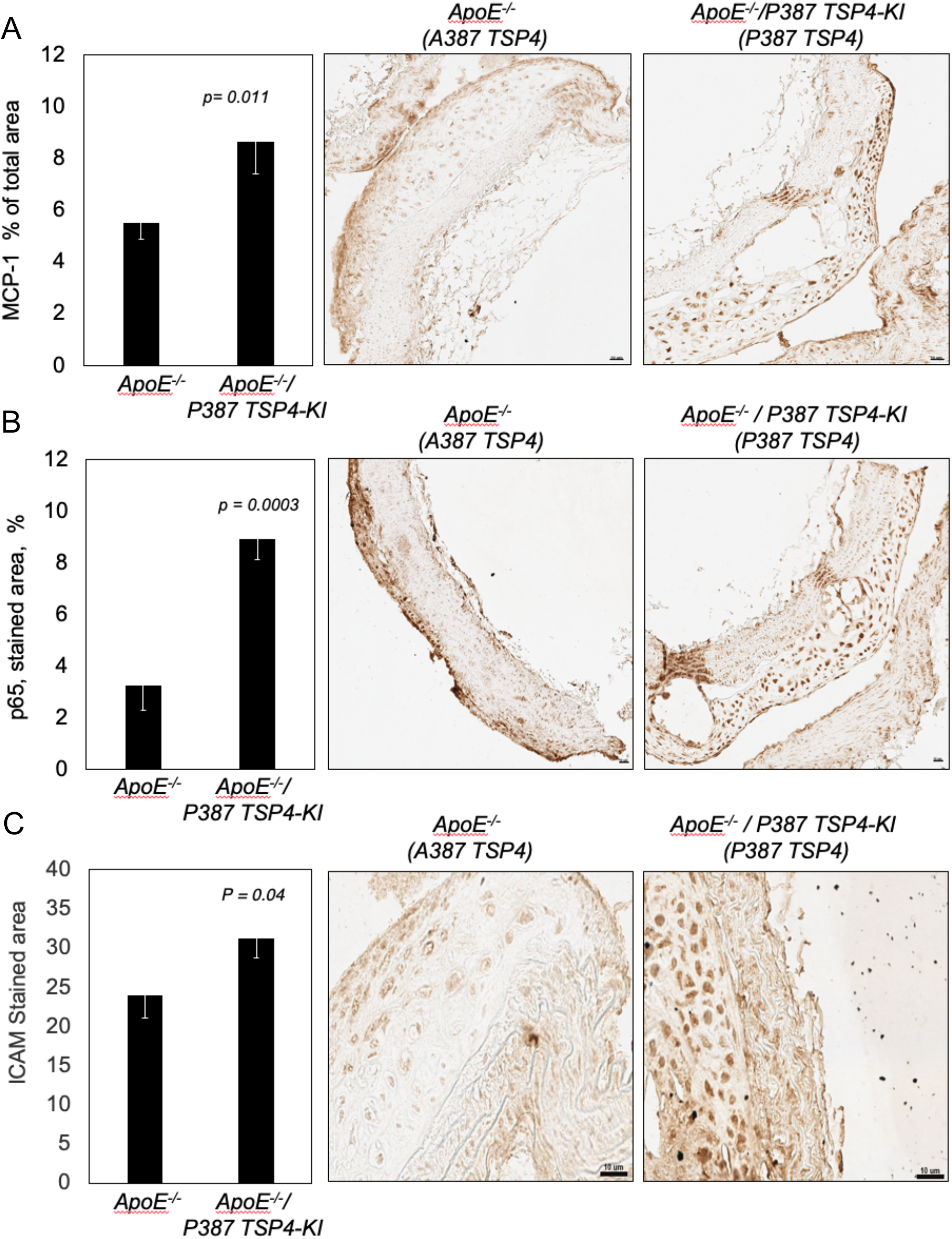
Increased levels of inflammation markers are associated with P387 TSP4. **A**:The level of MCP-1 is increased in the lesion of *ApoE*^*−/−*^/P387 TSP4 KI mice. MCP-1 staining in sections of atherosclerotic lesions in *ApoE*^*−/−*^ and *ApoE*^*−/−*^/P387-TSP4-KI mice aorta (n=10), immunohistochemistry with anti-MCP-1 antibody. **B:** The level of p65 is increased in the lesion of *ApoE*^*−/−*^/P387-TSP4-KI mice. p65 staining in sections of atherosclerotic lesions in *ApoE*^*−/−*^and *ApoE*^−/−^/P387 TSP4 KI mice aorta (n=10), immunohistochemistry with anti-p65 antibody. C: Increased levels of ICAM-1 in the lesion of *ApoE*^*−/−*^/P387-TSP4-KI mice. ICAM-1 staining in sections of atherosclerotic lesions in *ApoE*^*−/−*^and *ApoE*^−/xy2^/P387 TSP4 KI mice aorta (n=10), immunohistochemistry with anti-ICAM-1 antibody.

### 3.4. P387 TSP4 is associated with higher accumulation of macrophages in human coronary arteries

Left anterior descending coronary arteries were collected post-mortem after accidental death and genotyped to identify the vessels expressing P387 TSP4. Out of 23 collected vessels, 6 were heterozygous (HT). TSP4 and macrophages were visualized by immunofluorescence (Fig. 5) and immunohistochemistry (Fig. 6).

**Figure 5.**
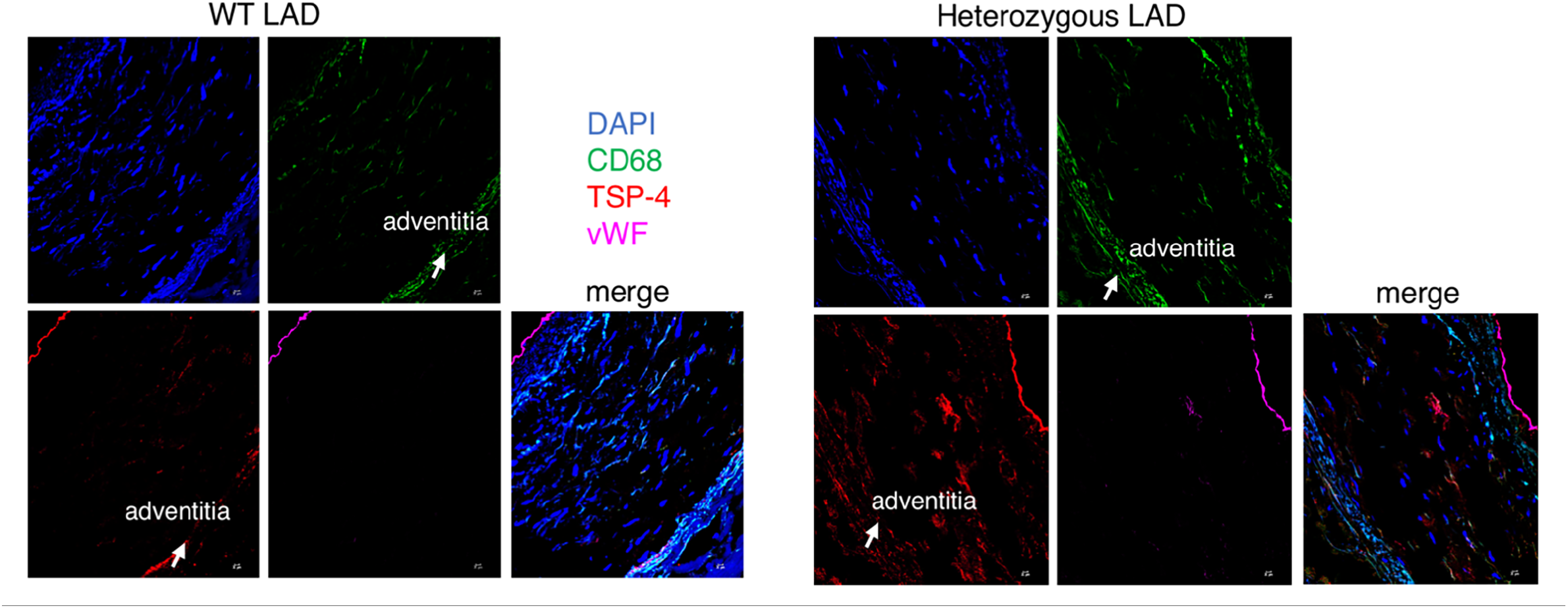
Macrophages in human atherosclerotic lesions of left anterior descending (LAD) coronary artery produce TSP4. **A:** LAD from A387 TSP4 carrier (WT LAD) and **B:** LAD from P387/A387 TSP4 carriers (Heterozygous LAD). Immunofluorescence of CD68, *Thbs4* and vWF, (TSP4 = red, anti–*Thbs4* Ab; CD68 = green, anti-CD68 Ab; vWF = magenta, anti-vWF Ab; nuclei = blue, DAPI).

**Figure 6.**
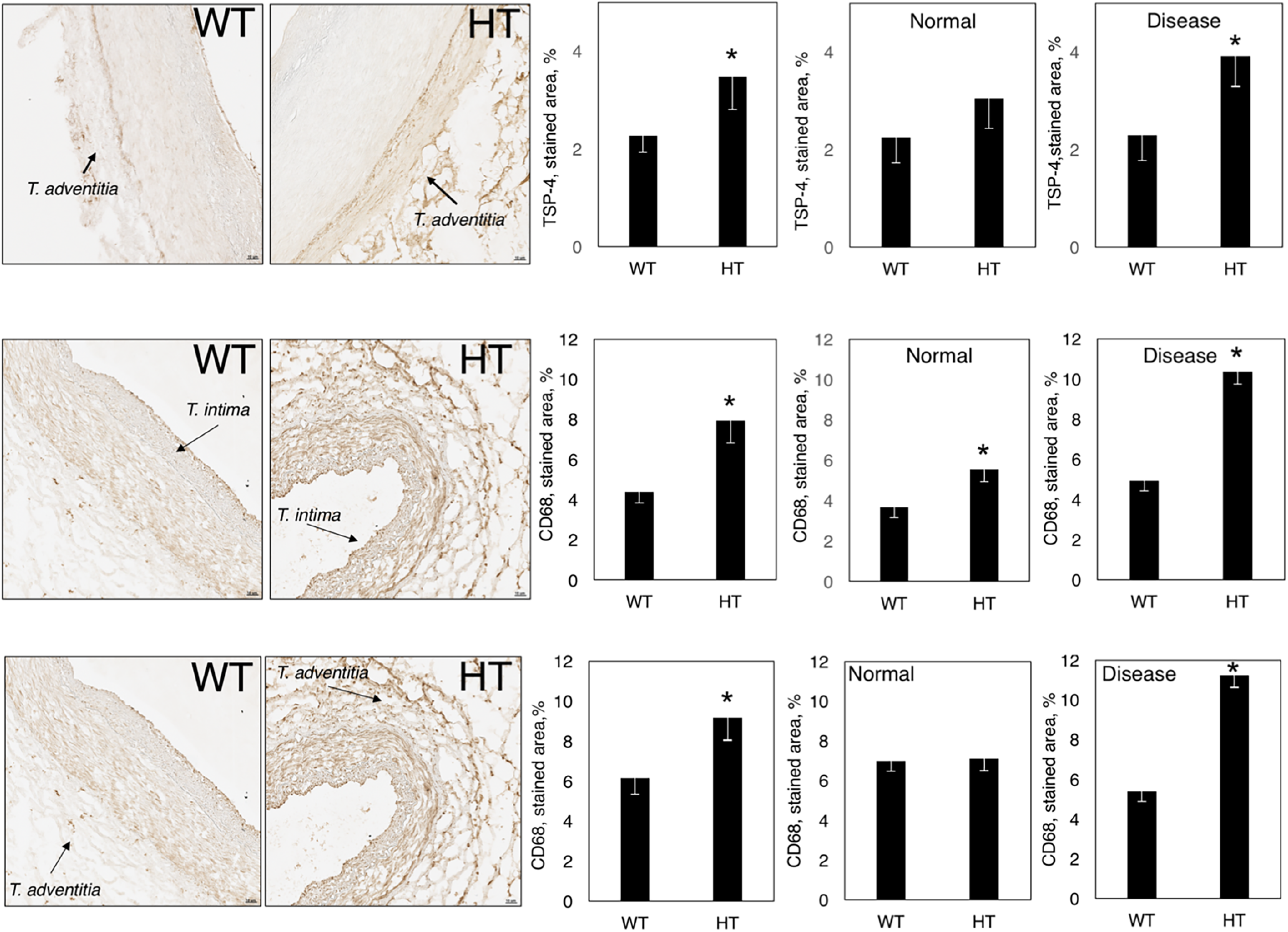
P387 TSP4 promotes accumulation of macrophages in human left anterior descending artery. **A:** Quantification of anti-*Thbs4* staining in all arteries (graph in the left), arteries without lesions (graph in the middle), and arteries with atherosclerotic lesions (graph in the right); WT (n=12) and heterozygous human arteries (HT, n=12). **B:** Macrophages (CD68) in *tunica intima* of human arteries. **C:** Macrophages (CD68) immunostaining in *tunica adventitia*.

In homozygous arteries expressing A387 TSP4 only, TSP4 was localized to endothelium and adventitia (Fig. 5A). In HT arteries, the levels of TSP4 were higher and easily detectable in all layers of the artery (Fig. 5B). In both genotypes, we detected the expression of TSP4 in macrophages (co-staining with anti-CD68 antibody), in addition to endothelial cells (EC, co-staining with anti-vWF antibody) and some vascular smooth muscle cells in *tunica media* (Fig. 5B).

TSP4 immunostaining was quantified in arteries. More TSP4 was detected in HT arteries, and higher levels were associated with atherosclerotic lesions than in the homozygous A387 TSP4 arteries (Fig. 6A). Macrophage accumulation correlated with TSP4 levels in the vessels: it was significantly increased in HT arteries, both in the *tunica intima* (Fig. 6B) and *tunica adventitia* (Fig. 6C), and was higher in atherosclerotic lesions (Fig. 6B and C).

### 3.5. P387 TSP4 is more active in supporting the adhesion of monocytes and macrophages

In seeking an underlying mechanism for the increased accumulation of macrophages with the P387 TSP4 variant, we compared the effects of A387 TSP4 and P387 TSP4 on monocyte/macrophage adhesion, migration, and activation using a macrophage-like cell line and mouse peritoneal macrophages.

The adhesion of human blood monocyte and the macrophage-like cells WBC264-9C was increased on recombinant P387 TSP4 (rTSP4) (Fig.7A and B). In the presence of P387 rTSP4, adhesion was associated with significantly higher H_2_O_2_ levels, even when corrected for differences in the number of adherent cells (Fig.7A), although both A387 and P387 TSP4 supported the adhesion of monocytes/macrophages and H_2_O_2_ production compared to wells coated with BSA.

**Figure 7.**
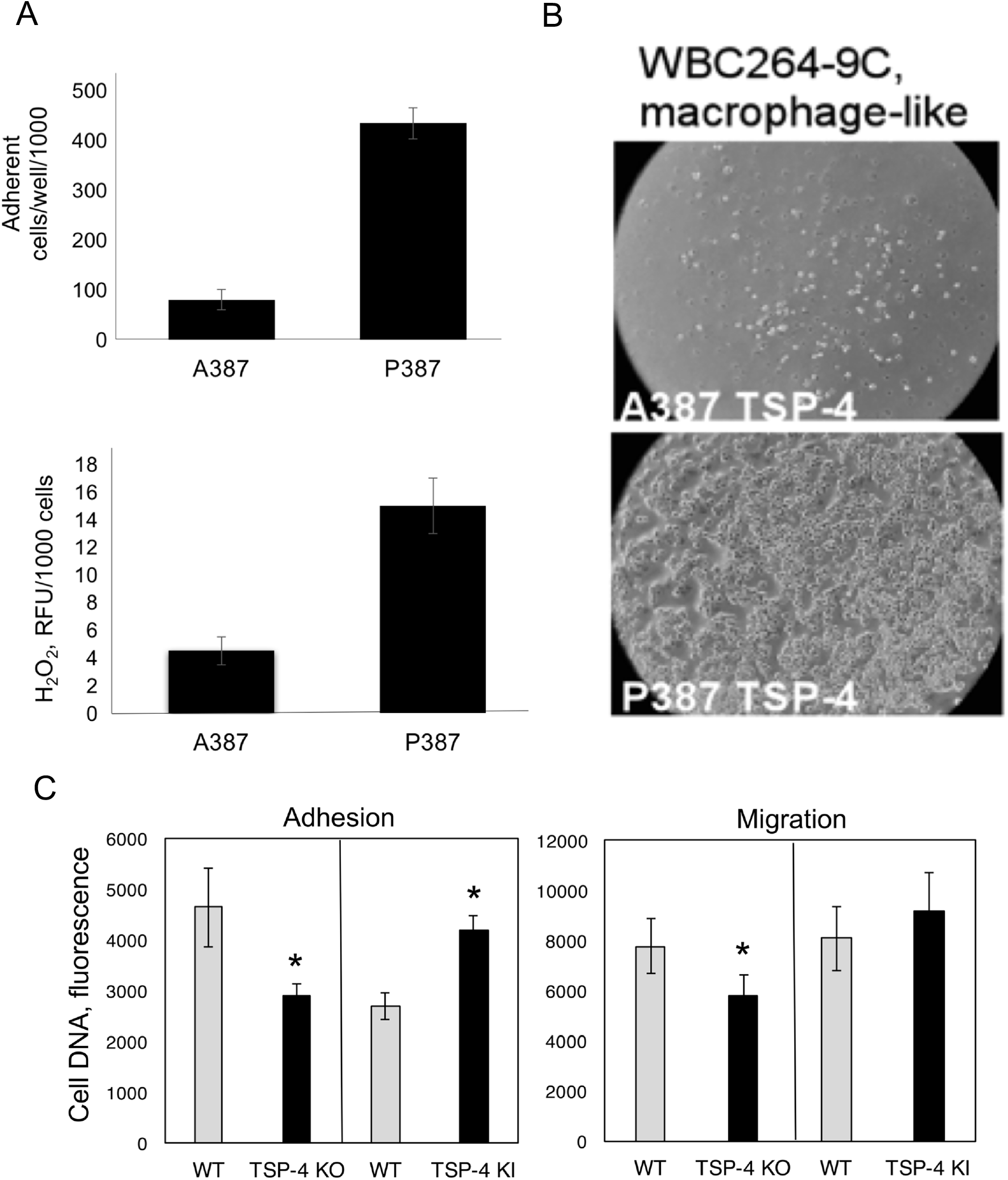
P387 TSP4 is more active in supporting adhesion of macrophages. **A**: Macrophage-like cells WBC264-9C and **B:** human monocytes purified from blood were plated in the cell culture dish coated with purified recombinant A387 TSP4 or P387 TSP4; n=3, p<0.05. C: Mouse peritoneal macrophages (MPM) from WT, *Thbs4*^−/−^, and P387-TSP4 KI mice treated with LPS (0.25 μg/g) for 72h were added to 24-well plates and incubated at 37°C for 1h (left, adhesion). Unattached cells were removed by washing, and the remaining DNA in the wells was measured using the CyQUANT reagent, n=12, *p<0.05. Right: Migration of MPM was measured Boyden chambers, 4h at 37°C, n=12, *p<0.05.

When peritoneal macrophages from WT, *Thbs4*^*−/−*^, and P387-TSP4-KI mice were used in adhesion and migration assays, both adhesion and migration were reduced in *Thbs4*^*−/−*^ mice compared to WT mice (Fig.7C). Cells from P387-TSP4-KI mice demonstrated increased adhesion, but did not have higher migratory capacity than cells from WT mice.

## 4. Discussion

In comparison to TSP1, the first discovered TSP (37) with a readily available source for purification from platelets, functions of TSP4 are poorly studied. However, recent reports stressing the importance of TSP4 in cardiovascular system (3–17), in cancer (15, 18–27), and in nervous system (28–32) have attracted a lot of attention to this matricellular protein. Although differences in the secondary and tertiary structure between TSP1 and TSP4 (1, 2, 38-41) suggested that their functions may be different too, only very recently have such differences been demonstrated experimentally. TSP4 was found to be pro-angiogenic (15, 27), unlike TSP1, which is a potent anti-angiogenic protein (42, 43). The role of TSP4 in inflammation was also reported to be different from the role of TSP-1. TSP1 promotes the resolution of inflammation in several animal models (44–46), while TSP4 was found to increase local vascular inflammation in atherosclerosis model. Specifically, atherosclerotic lesions in *ApoE*^*−/−*^ mice deficient in TSP4 were less cellular, contained fewer macrophages, and had reduced levels of inflammatory markers (17).

P387 SNP variant of TSP4 has been associated with increased risk of myocardial infarction and cardiovascular disease (3–10). We found that this variant of TSP4 is more active in its cellular interaction and effects on neutrophils and endothelial cells (14, 33–35). The effects of P387 TSP4 in atherosclerosis and its interaction with macrophages has not been previously characterized. Here, we examined the effect of P387 TSP4 on atherosclerotic lesions in *ApoE*^*−/−*^ mice and in human arteries from P387 carriers. A387 TSP4 increased early lesions in mice on chow diet, but was more inflammatory in lesion of mice on a Western diet, TSP4 increased macrophage accumulation without affecting the size of lipid lesions (17). Similar to WT A387 TSP4, P387 TSP4 did not change the size of atherosclerotic lesions visualized by Oil Red O staining. However, the cellularity of lesions was increased, and the immunostaining for CD68 revealed significant increase of macrophage accumulation in lesions of P387-TSP4-KI mice. The increased accumulation of macrophages in lesions supports our earlier observations of higher activity of P387 TSP4 compared to A387 TSP4 (14, 33–35).

P387 is a common SNP: initial study identified the prevalence of this variant at 34% in the US Caucasians population (6). P387 TSP4 variant acted as dominant and produced its CVD effects in heterozygous individuals. We collected and genotyped left anterior descending coronary arteries *post mortem* and identified a group of heterozygous (HT) P387 carriers. The localization of P387 TSP4 was similar to that of A387 TSP4 by immunofluorescence. We found that both variants accumulated in *tunica adventitia*, in luminal endothelial cells, and in *tunica intima* of atherosclerotic arteries. In both *t. adventitia* and *t. intima*, both variants of TSP4 were associated with macrophages, among other cells (e.g., smooth muscle cells and endothelial cells also known to produce TSP4)(16, 17, 35, 36). Quantification of macrophages in *t. adventitia* and *t. intima* using staining with macrophage marker CD68 revealed increased accumulation of macrophages in HT arteries within atherosclerotic lesions compared to homozygous A387 carriers or to the levels in arteries without lesions. The levels of P387 TSP4 in vessel wall were higher than the levels of A387 TSP4, probably reflecting the lower susceptibility of P387 TSP4 to proteolysis that we had noticed earlier (34). Furthermore, higher levels of P387 TSP4 allowed us to detect TSP4 production by macrophages. To date, production of TSP4 by the blood cells has not been reported. Additionally, quantification of markers of pro-inflammatory macrophages (CD38) and tissue repair macrophages (Egr2) revealed increased accumulation of pro-inflammatory macrophages associated with P387 TSP4. CD38 expression was associated with macrophages in lesions, while Egr2 was produced by both macrophages and other cell type, e.g., smooth muscle cells, which probably resulted in overestimation of tissue repair macrophages presence. Taken together with the previously reported effects of WT A387 TSP4 on vascular inflammation (17), these results suggested not only higher activity of P387 TSP4 in lesions but also the association of TSP4, especially P387 TSP4, with accumulation of pro-inflammatory macrophages in vascular tissue.

In addition to the increased number of the macrophages and the change in balance of pro-inflammatory and tissue-repair macrophage phenotypes, higher inflammation in the vessels of mice carrying P387 variant was demonstrated by increased levels of MCP-1, ICAM-1-1 and active p65 NFkB.

The observed association between TSP4 (17), especially P387 TSP4, and the increased accumulation of macrophages, suggested that the adhesive properties of these matricellular proteins might contribute to the enhanced macrophage content of atherosclerotic lesions. Earlier, we had reported increased adhesive activity of P387 TSP4 versus A387 TSP4 for neutrophils (33) and endothelial cells (14, 35). Using a macrophage-like cell line, primary mouse macrophages, and human monocytes, we found that TSP4 supported the adhesion of macrophages and monocytes. P387 was significantly more active in supporting adhesion of all cells tested. Increased adhesion was associated with higher release of H_2_O_2_ by human monocytes indicating increased activation. A387 TSP4 produced by macrophages supported both adhesion and migration as was demonstrated with peritoneal macrophages, but P387 produced by macrophages only affected adhesion without any effect on migration. The sources of TSPs are known to affect their effects, because the cellular effects depend on interactions of TSPs with other ECM ligands produced by cells. Cell-produced TSPs may exhibit different effects than exogenous purified recombinant TSP4. However, the supportive effects of TSP4, especially P387 TSP4, on adhesion have been consistent across all cell types tested.

Our results demonstrated that common P387 TSP4 variant accumulates in arteries in higher levels and is more active in promoting accumulation of pro-inflammatory macrophages and supporting inflammation in vascular wall. These observations provide a basis for the association of P387 TSP4 with cardiovascular disease and myocardial infarction and identify P387 variant as a SNP leading to enhanced inflammation in lesions and increased content of proinflammatory macrophages. Furthermore, the effects of TSP4 on macrophages in *ApoE*^*−/−*^ atherosclerosis and in human arteries with lesions set a precedence to propose its general role in chronic inflammation.

---

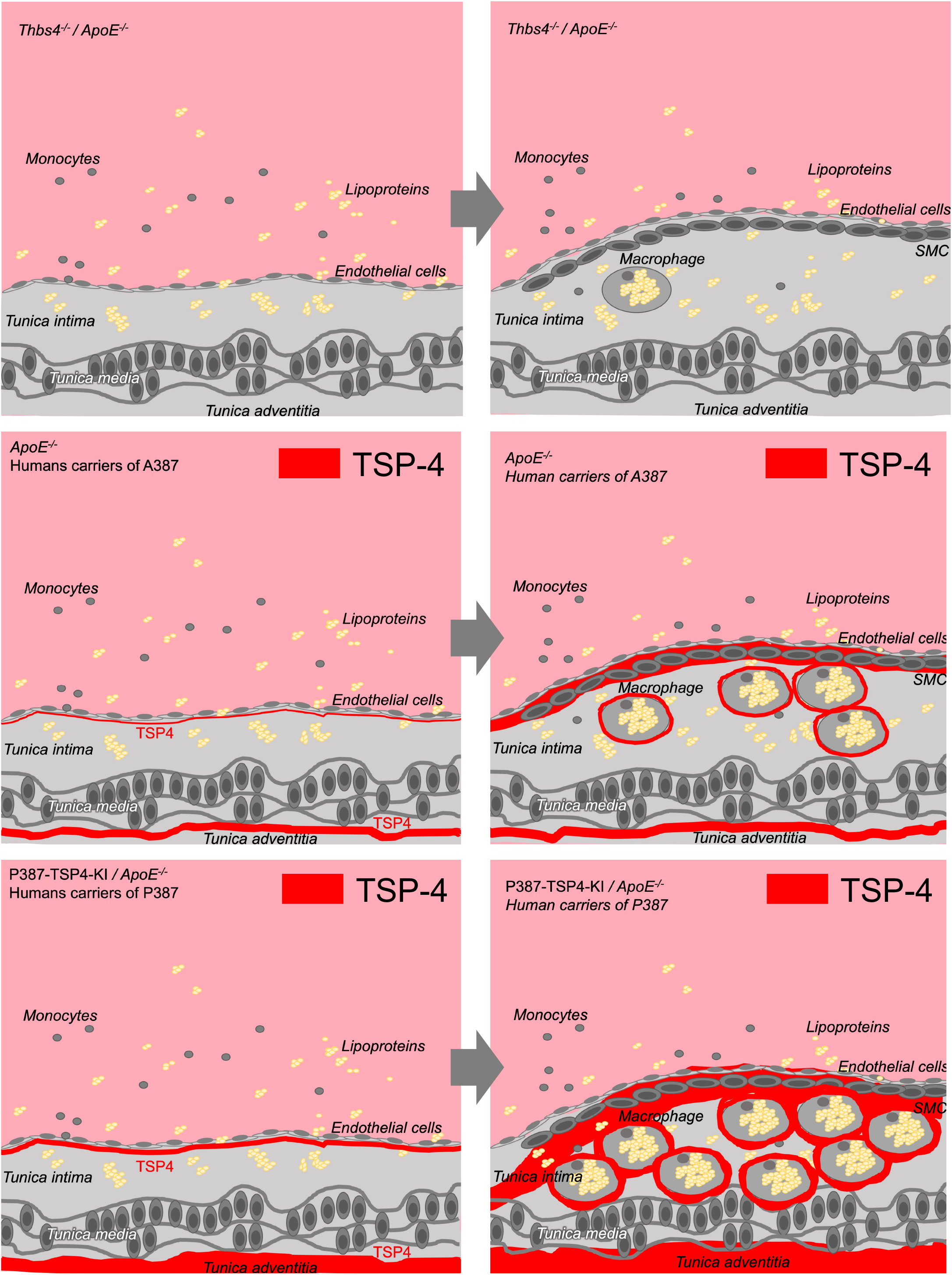

